# conn2res: A toolbox for connectome-based reservoir computing

**DOI:** 10.1101/2023.05.31.543092

**Authors:** Laura E. Suárez, Agoston Mihalik, Filip Milisav, Kenji Marshall, Mingze Li, Petra E. Vértes, Guillaume Lajoie, Bratislav Misic

**Affiliations:** McConnell Brain Imaging Centre, Montréal Neurological Institute, McGill University, Montréal, QC, Canada; Mila, Quebec Artificial Intelligence Institute, Montreal, QC, Canada; Department of Psychiatry, University of Cambridge, Cambridge, United Kingdom; Department of Mathematics and Statistics, Université de Montréal, Montreal, QC, Canada

## Abstract

The connection patterns of neural circuits form a complex network. How signaling in these circuits manifests as complex cognition and adaptive behaviour remains the central question in neuroscience. Concomitant advances in connectomics and artificial intelligence open fundamentally new opportunities to understand how connection patterns shape computational capacity in biological brain networks. Reservoir computing is a versatile paradigm that uses nonlinear dynamics of high-dimensional dynamical systems to perform computations and approximate cognitive functions. Here we present `conn2res`: an open-source Python toolbox for implementing biological neural networks as artificial neural networks. `conn2res` is modular, allowing arbitrary architectures and arbitrary dynamics to be imposed. The toolbox allows researchers to input connectomes reconstructed using multiple techniques, from tract tracing to noninvasive diffusion imaging, and to impose multiple dynamical systems, from simple spiking neurons to memristive dynamics. The versatility of the `conn2res` toolbox allows us to ask new questions at the confluence of neuroscience and artificial intelligence. By reconceptualizing function as computation, `conn2res` sets the stage for a more mechanistic understanding of structure-function relationships in brain networks.

## INTRODUCTION

Brains are complex networks of anatomically connected and functionally interacting neurons that have the ability to seamlessly assimilate and interact with a perpetually changing external environment [1]. Sensory stimuli elicit signalling events within structural connectivity networks and manifest as patterned neural activity. These emergent neural dynamics are thought to support the computations that underlie cognition and adaptive behaviour. However, a computational framework that describes how information processing and functional specialization occur in brain networks remains elusive. Developing such a framework would require understanding the multiple levels of the information-processing hierarchy, from how the brain’s network architecture shapes the complex activity patterns elicited by external stimuli, to how neural circuits extract from these evoked activity patterns the necessary information to compute with time-varying inputs.

How does network structure shape spatiotemporal patterns of neural activity, and how do neural dynamics support computations that underlie cognitive functions and behaviours? An important piece of the puzzle is the study of connectomics [2]. Technological and analytic advances in neuroimaging methods have made it possible to reconstruct the wiring patterns of nervous systems, yielding high-resolution connectomes of brains in multiple species [3–6]. The availability of connectomes has led to the formulation of a variety of models that aim to map network architecture to various functional aspects of the brain [7], such as emergent neural dynamics [8, 9], neural co-activation patterns [10] and inter-individual differences in behaviour [11–13] Multiple network features are correlated with emergent functional phenomena [14–19] but there is no clear mechanistic link between the static network architecture and cognition.

Furthermore, descriptive studies of the connectome across different species provide evidence that structural connectivity networks display topological features that are thought to shape the segregation and integration of information [20]. For instance, the simultaneous presence of a highly clustered architecture of segregated modules that promotes specialized information processing [21–27] and a densely interconnected core of high-degree hubs that shortens communication pathways and promotes the integration of information from distributed specialized domains [28, 29]. How these ubiquitous organization principles of the architecture of the brain confer computational capacity remains unknown.

Artificial intelligence offers alternative ways to approach the link between structure and function in brain networks that take into account computation [30, 31]. Within the expanding spectrum of artificial neural network models, reservoir computing makes it possible to describe how recurrent neural circuits extract information from a continuous stream of external stimuli and how they approximate complex time-varying functions [32, 33]. In reservoir networks learning occurs only at the readout connections, and hence the main architecture of the reservoir does not require specific weight calibration, remaining fixed throughout training. This eliminates a confounder while avoiding biologically implausible credit assignment problems such as the use of backpropagation training [34]. These reasons make reservoir computing an ideal paradigm to study the effects of connectome architecture on computation and learning. In this regard, machine-learning and artificial intelligence algorithms offer new ways to study structure-function relationships in brain networks by conceptualizing function as a computational property [31, 35].

Here we review the fundamentals of reservoir computing and how it can be applied to gain mechanistic insight about the information-processing of biological neural circuits. We then present `conn2res` (https://github.com/ estefanysuarez/`conn2res`.git), an open-source Python toolbox that implements connectomes as reservoirs to perform cognitive tasks. In the spirit of open-science, `conn2res` builds on top of and is interoperable with other third-party resources and research initiatives to offer a comprehensive corpus of tasks spanning a wide spectrum of computational and behavioral paradigms that researchers can experiment with. One of the hallmark features of the `conn2res` toolbox is the inclusion of multiple local neuron models. We have included a general use-case tutorial to illustrate the flexibility of the toolbox in terms of network architecture, local and global network dynamics, task paradigm, learning algorithm and performance metrics. While being inclusive of different modeling traditions, from micro-circuits to whole-brain network models, `conn2res` contributes a novel way for researchers to explore the link between structure and function in biological brain networks.

The present report has been conceived as a modular manuscript with 3 standalone components: (1) fundamentals of reservoir computing, (2) how reservoir computing has been applied to model brain phenomena and (3) an overview and tutorial of the `conn2res` toolbox. Readers who are new to the world of reservoir computing or curious how the paradigm can be applied to gain mechanistic insight into the information-processing properties of biological neural circuits can start with the review sections. Readers who are familiar with reservoir computing but wish to use the toolbox can skip ahead to the toolbox and tutorial sections.

### FUNDAMENTALS OF RESERVOIR COMPUTING

The conventional reservoir computing (RC) architecture consists of an input layer, followed by the reservoir and a readout module (Fig. 1) [32, 36, 37]. Typically, the reservoir is a recurrent neural network (RNN) of nonlinear units, while the readout module is a simple linear model. The readout module is trained to read the activation states of the reservoir — elicited by an external input signal — and map them to the desired target output in a supervised manner. In contrast to traditional artificial RNNs, the recurrent connections within the reservoir are fixed; only the connections between the reservoir and the readout module are learned (Fig. 1) [38].

**Figure 1.**
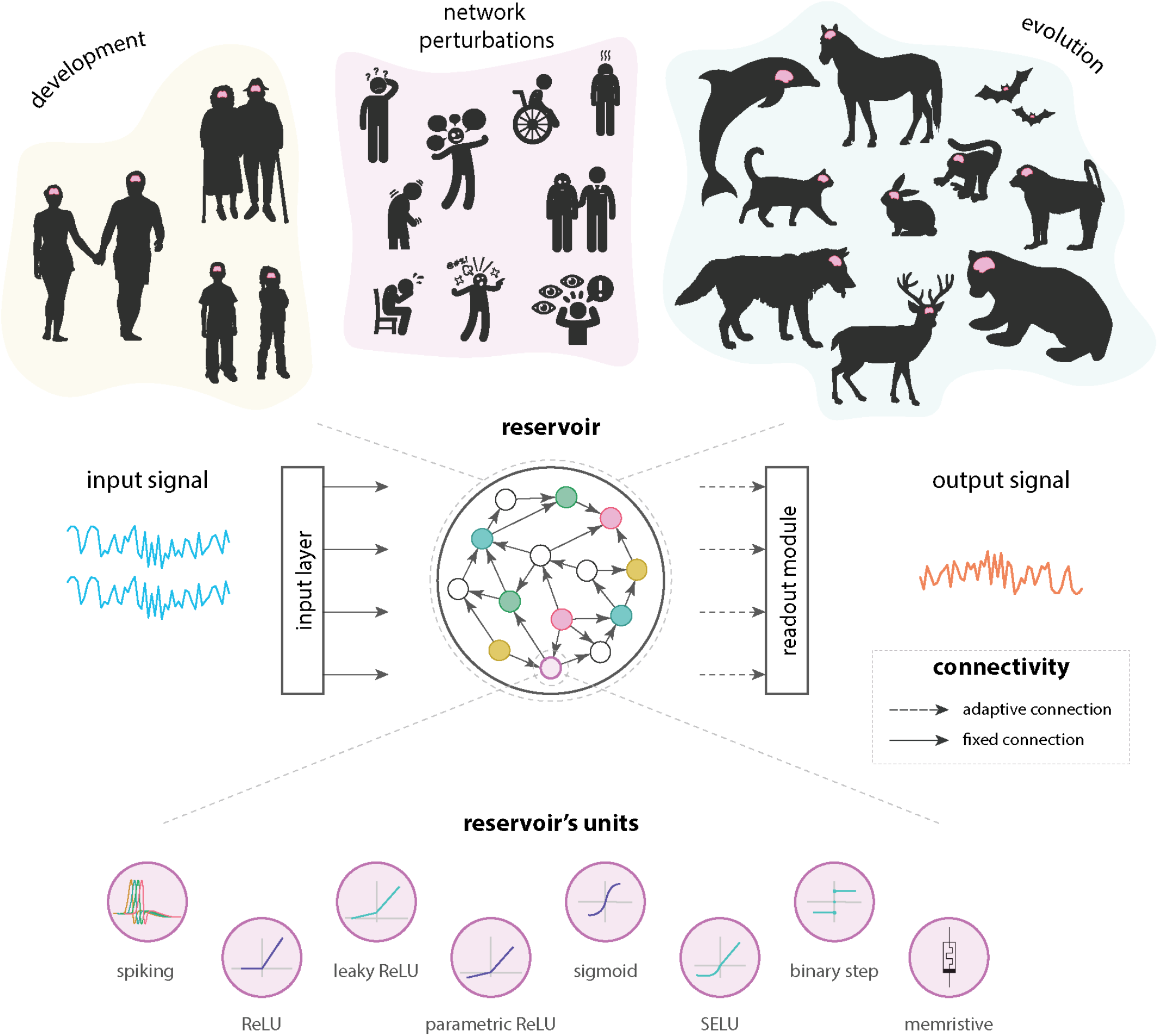
Reservoir computing. | The conventional reservoir computing architecture consists of an input layer, followed by a hidden layer, or *reservoir*, which is typically a recurrent neural network of nonlinear units, and the readout module, which is a simple linear model. In contrast to traditional artificial RNNs, the recurrent connections within the reservoir are fixed; only the connections between the reservoir and the readout module are trained. More importantly, RC allows arbitrary network architecture and network dynamics to be implemented by the experimenter. Hence, biologically-plausible wiring patterns (top panel) and different types of local dynamics (bottom panel) can be superimposed on the reservoir. In this way, the RC paradigm offers a tool for neuroscientists to investigate how network organization and neural dynamics interact to support learning in biologically-informed reservoirs.

So, how does RC work? RC capitalizes on the nonlinear response of high-dimensional dynamical systems, referred to as reservoirs. The reservoir performs a nonlinear projection of the input into a high-dimensional space. This transformation of the input converts nonlinearly separable signals into linearly separable ones such that a linear model in the readout module can be trained to map the transformed input to the desired output [38]. In other words, the reservoir converts inputs into rich dynamic patterns that contain integrated information about the history of inputs and are read out linearly to solve complex tasks. As long as the reservoir has sufficient built-in dynamical complexity and rich dynamics, a large variety of input-output mappings can be realized, including the approximation of complex timevarying functions, such as forecasting chaotic time series, considered to be a problem of high computational complexity. Under certain conditions, such as the presence of fading memory and separation properties, reservoirs can act as universal function approximators [32, 39–41]. The computational capabilities of the reservoir are determined by its dynamics, which arise from the interaction between the fixed architecture of the reservoir and the equations or rules governing the time evolution of its internal units. Importantly, unlike traditional artificial neural networks, in RC the network architecture can be set and the dynamics can be tuned by the experimenter. The RC paradigm thus offers the advantages that arbitrary architectures and dynamics can be superimposed on the reservoir, providing a tool for neuroscientists to investigate how connectome organization and neural dynamics interact to support learning in biologically-informed reservoirs.

The fact that RC can be used with arbitrary network architectures and dynamics, plus the possibility of performing a variety of tasks spanning multiple cognitive domains — from perceptual-motor functions, memory and learning, to complex attention and executive functions — makes it ideal to investigate how specific network attributes and dynamics influence the neural computations that support cognition [42]. Specifically, by implementing multiple tasks along these multiple cognitive domains, connectome-informed reservoirs allow us to systematically map network structure and dynamics to a unique set of identifiable computational properties exclusive to the task at hand. In this way, the RC framework allows us to build a comprehensive structural-functional ontology, relating network structure and dynamics to fundamental blocks of computation and, ultimately, to cognitive function.

The application of this hybrid approach between artificial intelligence and neuroscience goes beyond exploring the link between structure and function in the healthy brain. For instance, it can be applied in the clinical setting to study how neurological diseases affect learning in the network. By comparing the performance of reservoirs informed by clinical populations against those informed by healthy controls, this framework allows us to investigate whether cognitive decline, measured as variations in computational capacity, can be explained by measurable changes in network architecture due to neurodegeneration (Fig. 1). Another relevant application of the RC framework is the exploration of how the link between structure and function changes throughout adaptive processes such as development or evolution. For example, by implementing connectomes corresponding to different developmental phases, or species, as reservoirs, this framework allows us to investigate how variations in network architecture translate into differences in computational capacity across ontogeny and phylogeny (Fig. 1), respectively. In all cases, the RC framework allows for statistical significance testing by benchmarking empirical neural network architectures against random or null network models. The architectural flexibility of RC is multiscale: reservoirs can be informed by connectomes reconstructed at different spatial scales, from micro-circuits to meso- and macro-scale networks. Depending on the context, the units of the reservoir represent either populations of neurons or entire brain regions.

Compared to traditional RNNs, in which global network dynamics are determined by connectivity changes due to learning, the dynamical aspect of RC networks is equally flexible to their static structural counterpart. RC allows us to impose not only different types of local dynamics, but also global dynamics governing the population-level behaviour can be controlled. This means that the dynamical regime of the reservoir can be tuned to progressively transition from stable to chaotic dynamics, thus passing through a critical phase transition, or *criticality*. By parametrically tuning the dynamics to be closer or further from criticality, RC allows us to investigate the effects of qualitatively different neural trajectories near this critical point on the computational performance of the reservoir [43, 44]. An additional advantage of the dynamical and structural flexibility of RC is the possibility to enforce computational priors in the form of either functional or structural inductive biases [45]. Therefore, RC allows us to explore the functional consequences of information-processing strategies, such as critical dynamics or the presence of computational priors, thought to be exploited by biological brains [46–50].

Apart from the advantages that RC offers to the neuroscience community, this paradigm is also promising from an engineering point of view. Reservoirs can be realized using physical systems, substrates or devices, as opposed to – generally time- and energy-consuming — simulated RNNs [51, 52]. In this regard, the architecture of these neuromorphic chips could benefit from the emerging understanding of connection patterns in biological networks [53–55]. For instance, systematically mapping combinations of network attributes and dynamical regimes to a range of computational functions could assist the design of *ad hoc* or problem-specific tailored architectures. Due to their physical nature, neuromorphic systems are limited by spatial, material and energetic constraints, akin to biological neural networks. Because of this, insights gained about the economical organization of brain networks could contribute to the costeffective design of these information processing systems [35]. Furthermore, the fact that training only occurs at the readout stage makes RC an extraordinarily computationally efficient learning approach. In addition to this, parallel information-processing can be achieved by simultaneously training multiple readout modules to perform various parallel tasks. Therefore, physical RC and RC in general, provide a powerful method for faster and simpler multi-task learning, compared to other RNNs. Thanks to the dynamical and versatile nature of the reservoir, the RC paradigm is perfectly suited for a wide range of supervised tasks involving the processing of temporal and sequential data. These include: time series prediction, dynamical pattern generation, classification and segmentation, control, signal processing, and monitoring of rare events, among others [56]. Because of all these reasons, physical RC systems have become ideal candidates for the development of novel brain-inspired computing architectures [57].

### THE BRAIN AS A RESERVOIR

RC originated independently in the fields of machinelearning and computational neuroscience in the early 2000s with the seminal works of Herbert Jaeger on echo-state networks [36], and Wolfgang Maass on liquid state machines [32]. More broadly, the ideas encompassed by the RC paradigm had been around in different forms for more than two decades prior. A series of electrophysiological [58], behavioral [58, 63], and anatomical [59, 64–68] empirical findings in the late 1980’s and 1990’s gave rise to what would be later known as RC. Importantly, RC has been used to model biological mechanisms at multiple scales and here we briefly trace the origins and myriad applications of RC to different phenomena in neuroscience.

The first RC model was developed as a brain-inspired system that could explain physiological and behavioral phenomena observed in primates performing a sensorimotor sequence learning task [58, 60, 61]. This new model was built on a previous network model to understand saccadic movements in the primate corticostriatal oculomotor system [59]. The initial model, involving the posterior parietal cortex, frontal eye fields, superior colliculus and basal ganglia, was capable of reproducing the saccades observed in primates. To endow this circuit with the ability to display conditioned behavior in which the correct saccade target is selected among many, the prefrontal cortex was included and modeled as an RNN with fixed excitatory and inhibitory connections. Adaptive projections from the prefrontal cortex to the caudate were added and trained through reward-based learning. And so the first RC system was born: the prefrontal cortex — the *reservoir* — transformed visuospatial input from the posterior parietal cortex into activity sequences that were then associated with saccade sequences in the caudate — the *readout* — through dopamine-related reward learning (Fig. 2a). This circuit was able to reproduce the behavioral responses observed in a sensorimotor sequence learning task [60, 61].

**Figure 2.**
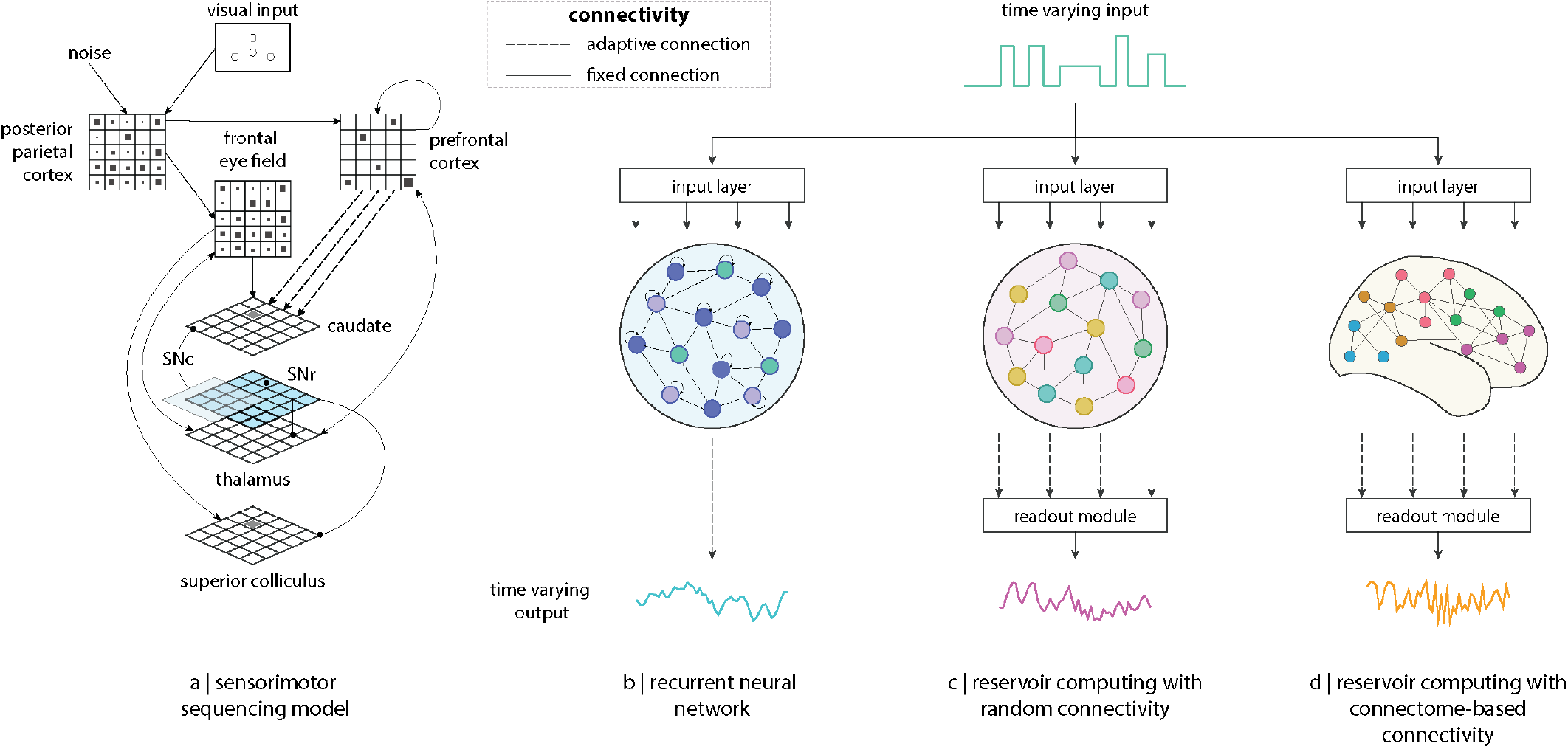
The evolution of reservoir computing. | (a) Sensorimotor sequencing model inspired by electrophysiological, anatomical and behavioral observations in primates [58, 59]. Here the prefrontal cortex is modeled as a recurrent neural network of excitatory and inhibitory neurons, and the caudate as a linear readout [60, 61]. Recurrent connections within the prefrontal cortex are kept fixed while connections from the prefrontal cortex to the caudate are learned via dopamine-related reward learning. This model is thought to be the first reservoir-like model ever implemented. (b) Generic recurrent neural network model. Different types of recurrent networks have been proposed to model computation in artificial and biological agents. The classic and most generic type of recurrent neural network used in artificial intelligence is characterized by the fact that recurrent connections are learned via backpropagation-through-time. Even though these networks are able to learn, biologically implausible network topologies emerge from training. On the other hand, random oscillatory recurrent neural networks with delayed coupling have often been used in neuroscience to model a variety of cortical phenomena, including the generation of temporally structured activity, or variable temporal relations among connected neurons used by hebbian-like plasticity mechanisms [62]. (c) In the classic reservoir computing architecture the reservoir consist of a recurrent neural network with randomly assigned weights. The topology of the reservoir remains fixed during training and learning occurs only at the connections between the recurrent network and the readout module. Examples of this include classic liquid state machines [32] and echo-state-networks [36]. (d) The advent of connectomics allows us to implement reservoirs with connectome-based architectures to explore the link between structure and function in brain networks from a computational point of view.

This new computational model proved to be versatile: it was sensitive to sequential and temporal structure [60, 61, 69, 70], and could also learn more abstract structures like arbitrary syntax rules [71, 72]. The model was also capable of higher cognitive functions such as language acquisition and processing [73], grammatical construction [74, 75], language production [76] and narrative [77], and even complex non-linguistic tasks such as exploration-exploitation [78] and optimized spatial navigation [79]. Remarkably, across all these different tasks, the activation of the units in the reservoir, and sometimes the readout, display a striking resemblance to what is observed in the living primate cortex under the same task contexts [78]. One such similarity is the activation of the reservoir’s units to complex nonlinear combinations of task dimensions, thus demonstrating a form of *mixed selectivity*, recently shown to be a computational strategy highly exploited by biological brains [80]. Altogether, these studies open an exciting possibility: the cortex could be conceived as, as Dominey himself put it, a set of “reservoirs organized in parallel interdigitated subsystems” [81].

A related notion of a reservoir as a delay-coupled recurrent oscillator network has been put forward as a conceptual framework to explain a variety of cortical phenomena (Fig. 2b) [62]. First, recurrently connected networks have the propensity to engage in oscillatory activity [82–85], generating temporally structured activity. This behaviour becomes useful in a variety of scenarios. For instance, it allows for pattern completion in the presence of partial information [86], supports the formation of novel associations [87], and permits the precise control of motor movements [88, 89]. Second, oscillatory activity can introduce precise but variable temporal relations between interconnected nodes in the network, including phase synchrony and phase shifts. Given the sensitivity of Hebbian-like learning mechanisms to temporal relations [90–93], the synchronization of oscillatory activity can favour the formation of synaptic connections and hence can be used for establishing associations. In this way, as posited by the “binding by synchrony” hypothesis [94–96], synchronous oscillations act as a binding mechanism for distributed nodes in the network that represent features of the same cognitive object, thus serving as a general code of relatedness [96]. Likewise, phase shifts can lead to rapid reversals of the direction of information flow, thus serving as gating mechanisms for information transfer according to the “communication through coherence” hypothesis [97]. Finally, imposing delays in recurrently connected networks gives rise to highly complex nonlinear dynamics that can be exploited for a variety of computational purposes. Low-dimensional stimuli are nonlinearly projected into a higher-dimensional space, converting nonlinearly separable stimuli into linearly separable. This allows the fast and effective classification of complex spatiotemporal input patterns [62]. Furthermore, the recurrent nature of the connections endows the network with a fading memory [62], that is, information about a stimulus persists for some time after the stimulus has disappeared. Therefore, information about multiple stimuli can be integrated over time, allowing for the encoding of temporal sequences [81]. Collectively these features showcase the versatility of the RC paradigm to capture the computational flexibility of the cortex.

Building on these early ideas, Wolfgang Maass proposed a related computational paradigm termed liquid computing [32, 98]. This framework was motivated by the question of how cortical columns in the brain seamlessly assimilate and process the spatial and temporal features of sensory stimuli [32, 98]. Maass and colleagues proposed a computational model whose construction was task-independent. The model comprised a recurrent network of spiking neurons in which only the connections of the readout neurons to the readout module are trained for the task at hand, while the recurrent connections within the network are fixed and randomly determined [32]. Connection probabilities are assigned based on the connectivity between laminae and neural populations [32]. These liquid state machines were shown to be successful for a variety of learning tasks involving the real-time processing of time-varying inputs [32]. Around the same time and independently, Herbert Jaeger developed echo-state networks [36], a similar framework conceived for artificial neural networks. Given their common goal of computing with nonlinear, high-dimensional dynamical systems, these two frameworks were brought together under the name of reservoir computing (Fig. 2c) [37].

More recently, advances in imaging and tracing technologies make it possible to reconstruct the wiring diagrams of multiple species, opening new opportunities to identify features of brain networks that theoretically enhance information-processing, such as modularity and small-worldness. Traditionally, reservoirs are randomly wired; by imposing some structure to this random connectivity we can explore how specific features of brain networks contribute to computational capacity (Fig. 2d). Modularity, for instance, affects information transmission in the network as well as its memory capacity [35, 99]. In random modular networks there is an optimal modularity where a balance between local cohesion and global communication is established, allowing the network to remember longer [99]. The same is true for empirical brain networks: the mesoscale modular organization of the human connectome, defined by intrinsic functional networks [100–105], and its underlying macroscale network topology, enhance the memory capacity of the reservoir, in particular when dynamics are critical [35]. Hence, in both random and empirical modular networks, there is an optimal level of modularity that delivers a balance between integration and segregation in the network [106], supporting optimal memory capacity [35, 99]. Similarly, small-world topology [21–23] — prevalent across biological networks [6] — contributes to the echo-state property of the reservoir [107], a measure of how flexibly network dynamics can untether from initial conditions, and promotes efficient signal propagation along the network [108]. Besides improving our understanding on how network architecture and dynamics affect the computational capacity of biological brain networks, implementing reservoirs informed by empirical connectomes can also be used for practical engineering purposes such as designing brainlike artificial neural network models or neuromorphic hardware [53, 109, 110].

Alternatively, the conventional RC architecture can be modified in unconventional ways to model different functional and structural aspects of the cortex. For example, a collaborative multimodal RC model can offer a mechanistic explanation to multisensory integration at early stages of the cortical processing hierarchy [111]. Multi-timescale changes in connectivity that occur through evolutionary and developmental processes can also be modeled using alternative learning paradigms, such as the Learning-to-Learn algorithm on modular and temporarily adaptive RC architectures [45]. These enforced structural priors — or pre-optimized reservoir’s hyper-parameters — facilitate subsequent learning in multi-task learning contexts [45, 112]. More recently, the concept of spatially-embedded recurrent neural networks was introduced [113]. These are recurrent networks with adaptive weights, confined within a 3D Euclidean space, and whose learning is constrained by biological optimization processes, like the minimization of wiring costs or the optimization of interregional communicability, in addition to the maximization of computational performance. When the pruning of the network is guided by these biological optimization principles, the resulting network architecture displays characteristic features of biological brain networks, including a modular structure with a small-world topology, and the emergence of functionally specialized regions that are spatially co-localized and implement an energetically-efficient, mixed-selective code [80, 113]. Altogether, reservoir computing, and recurrent neural network models in general, open a world of possibilities to provide mechanistic explanations for the way computations take place in real brain networks. By avoiding biologically implausible credit assignments derived from brackpropagation training [34], reservoir computing becomes ideal for this purpose.

### THE conn2res TOOLBOX

`conn2res` is an open-source Python toolbox that allows users to implement biological neural networks as reservoirs to perform cognitive tasks (https://github.com/estefanysuarez/conn2res.git). The tool-box is built on top of the following well established, documented and maintained Python package dependencies: `NumPy` (https://numpy.org; [114–116]), SciPy (https://scipy.org; [117]), `pandas` (https://pandas.pydata.org; [118]), `Scikit-Learn` (https://scikit-learn.org; [119]), `Gym` (https://www.gymlibrary.dev; [120]), `NeuroGym` (https://neurogym.github.io; [42]), `bctpy` (https://github.com/aestrivex/bctpy; [121]), `Seaborn` (https://seaborn.pydata.org; [122]) and `Matplotlib` (https://matplotlib.org; [123]). The toolbox is also interoperable with other relevant Python packages including `The Virtual Brain` (https://www.thevirtualbrain.org; [124]), `bio2art` (https://github.com/AlGoulas/bio2art; [109]), `NeuroSynth` (https://neurosynth.org; [125]), `Neuromaps` (https://neuromaps-main.readthedocs.io; [126]) and the `Enigma Toolbox` (https://enigma-toolbox.readthedocs.io; [127]).

Besides its extensive interoperability with other Python packages, a major strength of the `conn2res` tool-box is its flexibility in the choice of the different components that are part of the main RC workflow, in particular, the possibility to implement arbitrary network architectures and different types of local dynamics, which to our knowledge are typically fixed in other RC packages. The baseline `conn2res` workflow requires the following input arguments (Fig. 3a): **(*i*) task name or dataset:** the name of the task to be performed, or a labeled dataset of input-target pairs for supervised learning can also be provided. `conn2res` is a wrapper of `NeuroGym` [42], a curated collection of behavioural paradigms that were designed to facilitate the training of neural network models, and that are relevant for the neuroscience community. All of the 20+ tasks available in `NeuroGym` are also available in `conn2res` — some of these include perceptual decision-making, context-dependent decision making, delayed comparison, delayed-paired association and delayed match category —; **(*ii*) connectome:** the connectivity matrix, which serves as the reservoir’s network architecture. The toolbox supports binary and weighted connectivity matrices of both directed and undirected networks; **(*iii*) input nodes:** the set of nodes that receive the external input signals concerning the task at hand; **(*iv*) readout nodes:** the set of nodes or modules from which information will be retrieved to train the linear model in the readout module; **(*v*) reservoir local dynamics:** the type of dynamics governing the activation of the reservoir’s units. Right now only discrete-time, linear and nonlinear dynamics can be implemented. These include artificial neuron models with different activation functions such as ReLU, leaky ReLU, sigmoid or hyperbolic tangent. In the future, continuous-time nonlinear dynamical models will be implemented, including spiking neuron models; and lastly the type of **(*vi*) linear model:** specified as an instance of a linear model estimator from the `Scikit-Learn` library to be implemented for learning by the readout module [119].

**Figure 3.**
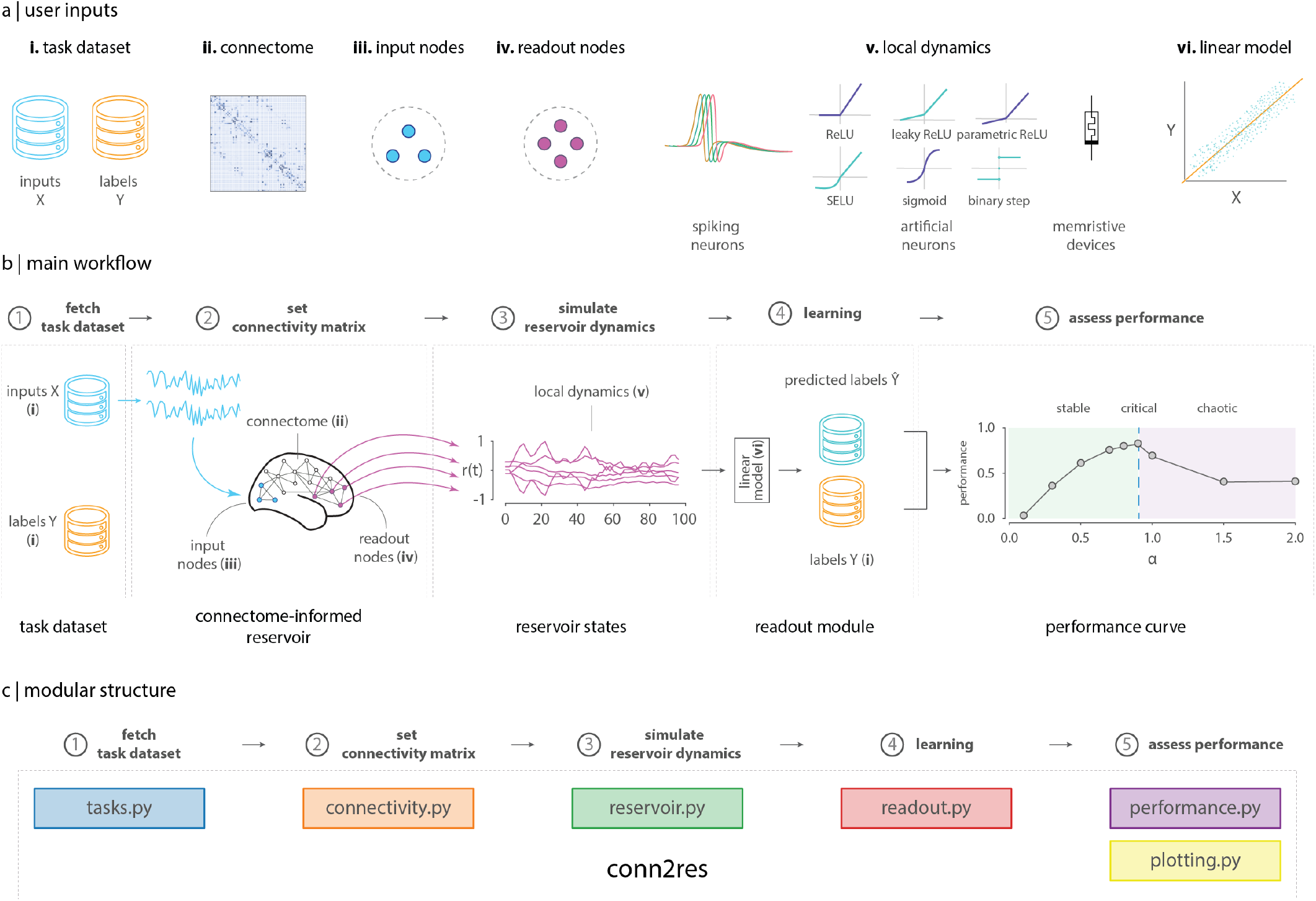
conn2res toolbox. | (a) The general `conn2res` workflow requires the following parameters to be provided by the user: *i*) a task name or a supervised learning dataset; *ii*) a connectome or connectivity matrix; *iii*) a set of input nodes; *iv*) a set of readout nodes or modules; *v*) the type of local dynamics, which can be either spiking neurons, artificial neurons (with a variety of activation functions), or memristive devices (for the simulation of physical reservoirs); and *vi*) the linear model to be trained in the readout module. (b) In the mainstream `conn2res` workflow the input signal (X) is introduced to the reservoir through the input nodes (blue nodes). The signal propagates through the network, activating the states of the units within the reservoir. The activation states of the readout nodes (purple nodes) are then retrieved and used to train a linear model to approximate the target signal (Y). Depending on the type of the reservoir, the performance can be a single score or a curve of performance as a function of a parameter that tunes the dynamics of the reservoir. (c) The `conn2res` toolbox has a modular architecture. It consists of six modules, each one comprising functions that support a specific step along the `conn2res` pipeline.

The typical `conn2res` workflow is depicted in Fig. 3b. In the first stage, **fetch task dataset**, a supervised dataset consisting of input-label pairs is either fetched from the `conn2res` repository, if the name of the task is provided by the user, or directly loaded if an external path is provided instead. In the second stage, **set connectivity matrix**, an instance of a reservoir object is created, and its network architecture and dynamics are set based on the connectivity matrix and the type of local dynamics specified by the user, respectively. In the third stage, **simulate reservoir dynamics**, the task inputs from the previous stage are introduced as external signals to the reservoir through the set of input nodes specified by the user. The dynamical models in the `conn2res` toolbox simulate the time evolution of the reservoir’s internal units (which are activated thanks to the propagation of the external input signals), and generate time series activity for each node in the reservoir. In the fourth stage, **learning**, the time series activity of the readout nodes or modules specified by the user are retrieved and passed on to the readout module, together with the task labels from the first stage. Both of these are used to train the linear model in the readout module. Finally, during the fifth and last stage, **assess performance**, depending on the nature of the reservoir, the final output can be either a single performance score, or a performance curve that displays performance as a function of the parameter that controls for the qualitative behavior of the reservoir’s dynamics (i.e., stable, critical or chaotic). Various performance metrics are currently available depending on whether the task requires a classification or a regression model. To facilitate the user’s experience, the toolbox provides several example scripts that illustrate use-case driven workflows.

The `conn2res` toolbox has a modular design. It consists of six modules, each one containing functions that support a specific step along the mainstream `conn2res` pipeline (Fig. 3c). The wrapper functions and classes used to generate the task datasets can be found in the ***tasks*.*py*** module. All types of manipulations on the connectivity matrix, such as binarization, weight scaling, normalization and rewiring, are handled by the Conn class in the ***connectivity*.*py*** module. All the reservoir’s features including its network architecture, local dynamics and retrieval of reservoir’s activation states, are handled by the Reservoir class in the ***reservoir*.*py*** module. The functions in charge of the training and test of the linear model in the readout module are contained in the ***readout*.*py*** and ***performance*.*py*** modules, respectively. Finally, the ***plotting*.*py*** module offers a set of plotting tools that assist with the visualization of the different types of data generated along the pipeline, including the task input-output data, the 2D connectivity matrix of the reservoir’s network architecture, the simulated reservoir states, the decision function of the readout module, and the performance curve.

### TUTORIAL

In this section we present a toy example in which we use `conn2res` to evaluate the effect of different types of local and global dynamics on the performance of a connectome-informed reservoir across two cognitive tasks: perceptual decision making [128] and contextdependent decision making [129]. To do so, we implement an echo-state network [36] whose connections are constrained by a human connectome reconstructed from diffusion-weighted MRI data. To select the set of input and readout nodes, we use a functional connectivity-based partition of the connectome into intrinsic networks [104]. We define input nodes as a set of randomly selected brain regions from the visual system, and for the readout nodes we select all brain regions in the somatomotor system. Local dynamics are determined by the activation function of the reservoir’s units. Here we use sigmoid and hyperbolic tangent activation functions. Global network dynamics are set by parametrically tuning *α*, which corresponds to the spectral radius of the connectivity matrix [130]. The dynamics of the reservoir are considered to be stable if *α <* 1, and chaotic if *α >* 1. When *α ≈* 1, the dynamics are said to be critical [46]. Because both tasks can be treated as supervised classification problems, we use a Ridge classifier model to train the readout module. We generate 1000 trials per task (70% training, 30% test), and we perform each task using 50 different realizations of the task’s labeled dataset. The distribution of performance scores is reported across the 50 instances of the task dataset.

Next we walk the reader through each of the steps along the main `conn2res` pipeline, and use the visualization tools included in the ***plotting*.*py*** module to depict the main output at each stage, facilitating the conceptual understanding of the workflow. Details about the practical implementation can be found in the *examples* folder of the `conn2res` toolbox. Results for the perceptual and context-dependent decision making tasks are shown on the left and right columns of Fig. 4, respectively. The first top panel in Fig. 4 consists of a single plot that displays the time series of the input (*x*_*i*_) and target labels (*y*) obtained during the task dataset fetching process. The perceptual decision making task is a two-alternative forced choice task in which the reservoir must be able to integrate two stimuli to decide which one is higher on average (left column in Fig. 4). In the context-dependent decision making task the reservoir has to perform one of two different perceptual discriminations, indicated by a contextual cue in every trial (right column in Fig. 4). Trials are delimited by vertical black dotted lines.

**Figure 4.**
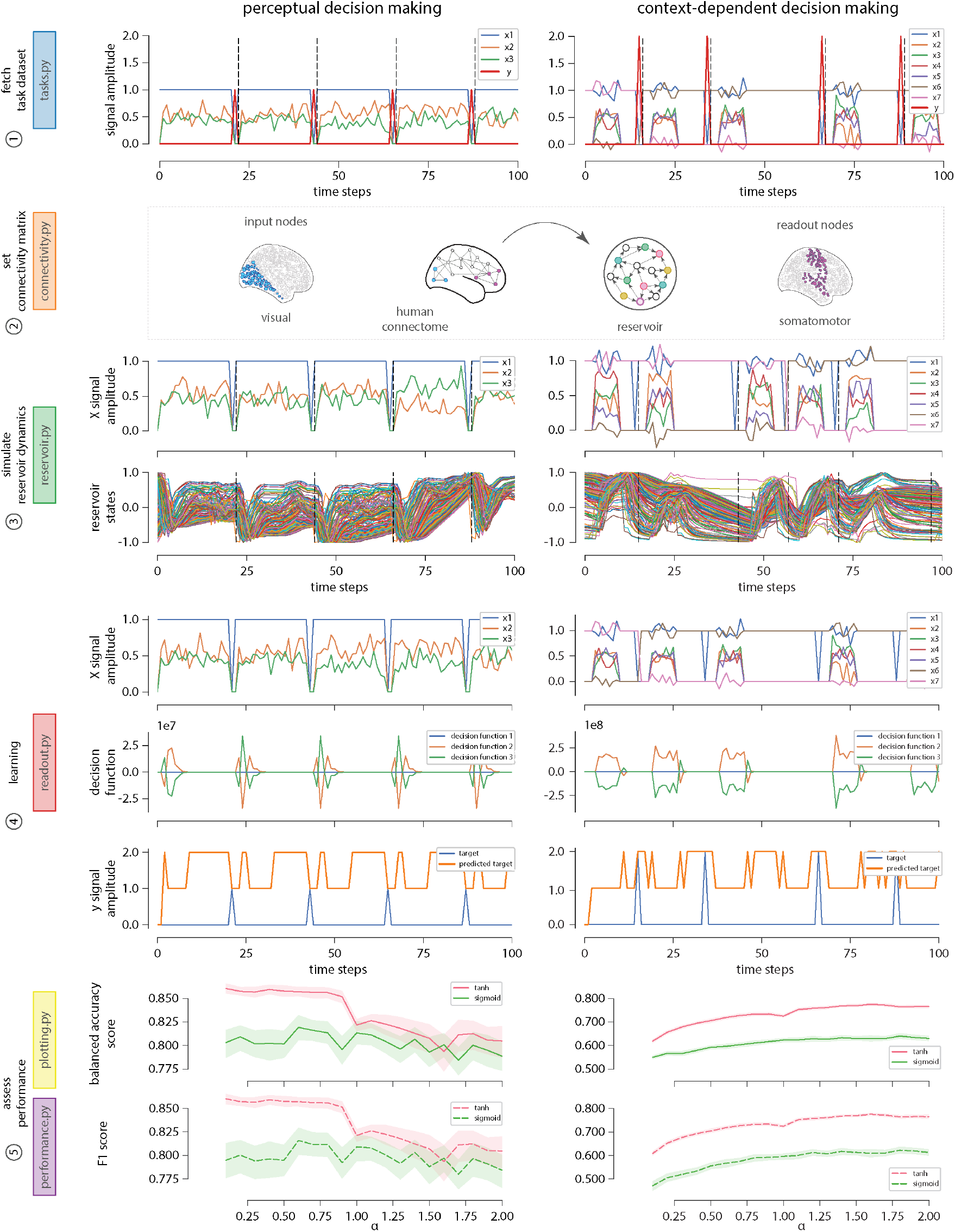
Use-case tutorial: effect of local and global dynamics on the performance of perceptual and context-dependent decision making tasks. | Perceptual decision making (left column): is a two-alternative forced choice task (*y* = {1, 2}) in which the reservoir must be able to integrate two stimuli (*x*_2_ and *x*_3_; *x*_1_ serves as a bias) to decide which one is higher on average. Context-dependent decision making task (right column): in this task the reservoir has to perform one of two different perceptual discriminations (*y* = {1, 2}), indicated by a contextual cue in every trial (determined by *x*_1_ to *x*_7_). From top to bottom: the first panel displays the time series of the input (*x*_*i*_) and target (*y*) signals obtained during the task dataset fetching step. The second panel presents a toy representation of the assignment of a connectome-based connectivity matrix to the reservoir’s network (center). It also shows the set of input (left) and readout (right) nodes selected for the analysis. The third panel displays the simulated reservoir’s dynamics; the top plot shows the time series of the input signals and the bottom plot shows the simultaneous activation states of the readout nodes within the reservoir (results shown here correspond to the simulated reservoir states with hyperbolic tangent as activation function). The fourth panel illustrates the learning process that takes place in the readout module during training. At every time step, the top plot shows the input signals, the middle plot shows the decision function of the classifier — in the readout module —, and the bottom plot shows the predicted versus the target signal. Finally, the fifth panel shows the performance curves as a function of both local (hyperbolic tangent in pink and sigmoid in green) and global (varying *α*) network dynamics. Two metrics were used to measure the performance of the classification: balanced accuracy (top - solid line) and F1 score (bottom - dotted line).

The second panel in Fig. 4 from top to bottom is a toy representation of the assignment of the connectome-based connectivity matrix to the reservoir’s network (center); it also shows the input nodes (left) used for the introduction of the external input signal into the reservoir during the simulation of the reservoir’s dynamics, and the readout nodes (right) used for the retrieval of information from the reservoir during the learning phase. The third panel in Fig. 4 depicts the simulation of the reservoir’s dynamics and it consists of two plots: the top plot presents the time series of the input signals (*x*), while the bottom plot shows the simultaneous reservoir’s activation state at every time step. These plots help the user visualize how reservoir states evolve as a function of the external inputs. The fourth panel in Fig. 4 makes reference to the learning process of the readout module during training. This panel contains three plots: the time series of the input signal (top), the decision function of the Ridge classifier (middle), and the predicted versus the ground truth target signals (bottom). Finally, the fifth panel in Fig. 4 shows the performance of the reservoir as a function of both local and global network dynamics. This panel presents two plots, each one corresponding to a different classification performance metric: balanced accuracy (top) and F1 score (bottom). Each plot displays two curves that indicate how performance varies as a function of *α*, and each curve corresponds to a different activation function: hyperbolic tangent (pink) and sigmoid (green).

Results in Fig. 4 suggest that both local and global network dynamics have an impact on task performance. At the local level, both tasks, perceptual and contextdependent decision making, benefit from having a hyperbolic tangent activation function, compared to the sigmoid. However, dependence of task performance on global network dynamics varies from one task to the other. Computations required in the perceptual decision making task take advantage of stable network dynamics if the local nonlinearity is hyperbolic tangent, whereas with a sigmoid nonlinearity the network does not show a strong dependence with respect to global network dynamics. In contrast, performance in the contextdependent decision making task improves as global network dynamics transition from stable to chaotic, regardless of the type of local activation function. As expected, the effect of local and global network dynamics on task performance depends on the type of computations required by the task at hand.

This example helps us to illustrate the flexibility of the `conn2res` toolbox in terms of the choice of network architecture, local and global network dynamics, computational properties, performance metrics and learning algorithms. Even though the type of experiments that the `conn2res` toolbox has been designed for have more of an exploratory character, we expect that as imaging technologies improve together with our understanding of the anatomical structure of biological brains, more hypothesis-driven type of experiments can be carried out with `conn2res`.

## CONCLUDING REMARKS

Despite common roots, modern neuroscience and artificial intelligence have followed diverging paths. The advent of high-resolution connectomics and the incredible progress of artificial neural networks in recent years present fundamentally new and exciting opportunities for the convergence of these vibrant and fast-paced fields. Here we briefly summarized the principles of the RC paradigm, reviewed various ways in which it can be applied to model diverse cortical phenomena, and introduced `conn2res`, an open-source code initiative designed to promote cross-pollination of ideas and bridge multiple disciplines, including neuroscience, psychology, engineering, artificial intelligence, physics and dynamical systems. Below we look outward and propose how the `conn2res` toolbox can address emerging questions in these fields.

The `conn2res` toolbox embodies the versatility of the RC paradigm itself. By allowing arbitrary network architecture and dynamics to be superimposed on the reservoir, `conn2res` can be applied to investigate a wide range of neuroscience problems: from understanding the link between structure and function, studying individual differences in behaviour, to exploring the functional consequences of network perturbations, such as disease or stimulation, or the computational benefits of specific architectural features, such as hierarchies and modules. The `conn2res` toolbox can readily accommodate network reconstructions at different spatial scales, from microcircuits to large-scale brain networks, and obtained using different imaging modalities, such as tract-tracing or diffusion MRI. Networks reconstructed at different points in either development and evolution can also be implemented in the toolbox to study, for instance, how structural adaptations across ontogeny and phylogeny shape computational capacity in brain networks. Collectively, `conn2res` offers new and numerous possibilities to discover how computation and functional specialization emerge from the brain’s anatomical network structure.

RC is often presented as a unified framework to train RNNs, but in a broader sense, it is a general framework to compute with high-dimensional, nonlinear dynamical systems, regardless of the choice of reservoir! Since any high-dimensional physical system with nonlinear dynamics could serve as reservoir — and these are abundant in both natural and man-made systems — a new field of applied research has emerged: physical reservoir computing. Here the goal is to exploit the rich dynamics of complex physical systems as information-processing devices. Physical substrates used for reservoirs are quite diverse: from analog circuits [131–134], field programmable gate arrays [135–138], photonic opto-electronic devices [139–144], spintronics [145–147], quantum dynamics [148, 149], nanomaterials [150–156], biological materials and organoids [157– 162], mechanics and robotics [163–165], up to liquids or fluids [166, 167], and most recently, origami structures [168]. As physical reservoir computing becomes more popular, we envision the use of `conn2res` as a workbench to explore the effect of network interactions on the computational properties of physical reservoirs. Anticipating this, `conn2res` is currently equipped with a dedicated class for physical reservoirs, which allows memristive networks — a promising alternative for neuromorphic computing [152] — to be implemented as reservoirs. In this sense, the paradigm and the `conn2res` toolbox can be applicable to a wide variety of problems in adjacent scientific disciplines. From the neuro-connectomics perspective, `conn2res` offers new and numerous possibilities to discover how structure and function are linked in biological brain networks. From the artificial intelligence perspective, reverse-engineering biological networks will provide insights and novel design principles for re-engineering artificial, brain-inspired RC architectures and systems.

Altogether, `conn2res` is an easy-to-use toolbox that al-] lows biological neural networks to be implemented as artificial neural networks. By combining connectomics and AI, the RC paradigm allows us to address new questions in a variety of scales of description and many adjacent fields. We hope that by reconceptualizing function as computation, `conn2res` allows us to take the next step towards understanding structure-function relationships in brain networks.

